# The metacognitive signatures of imposterism

**DOI:** 10.64898/2026.11.22.767833

**Authors:** Greta Mohr, Florencia Imakulata, Robin A. A. Ince, Christopher S.Y. Benwell

## Abstract

The imposter phenomenon (or imposterism) is a commonly reported psychological pattern in which individuals persistently doubt their abilities and accomplishments despite objective evidence of their competence. Imposterism is associated with chronic self-doubt and reduced well-being, yet its cognitive mechanisms remain unclear. In a large online sample (N = 602), we investigated how self-reported imposter feelings relate to decision-making, confidence and information search in a perceptual decision-making task. Imposterism was associated with lower confidence in perceptual decisions and increased information seeking, despite not being associated with any detriment in objective perceptual performance. Though symptoms of anxiety and depression contributed to the relationship between imposterism and confidence, imposterism uniquely predicted increased information-seeking.

Interestingly, the increased information seeking associated with imposterism did not lead to an imposterism-specific increase in final decision performance and did not alleviate overall uncertainty: imposterism remained associated with reduced confidence even for final decisions. To explain this, we hypothesised that information seeking may compensate for maladaptive second-guessing. Supporting this, imposterism predicted greater response switching and decreases in accuracy on trials where no additional information was sought, reflecting counterproductive changes of mind. These effects were absent on trials where information was accessed, suggesting that seeking additional information helped buffer against maladaptive second-guessing.

Overall, our findings show that imposterism is marked by low confidence, heightened information seeking, and vulnerability to second-guessing in the absence of external reassurance. These results provide novel insights into distortions of self-evaluation in imposterism and highlight a compensatory role for information seeking in stabilising performance. Our study advances understanding of a widely discussed yet underexplored psychological phenomenon with significant implications for education, workplace wellbeing, and mental health.

## Introduction

The imposter phenomenon (or imposterism) involves feelings of inadequacy and self-doubt with respect to one’s intellectual and professional abilities, despite evidence of competence and success, which result in fear of being exposed as a ‘fraud’ or ‘imposter’ ^[1]^. These feelings are posited to stem from an inability to internalise achievements, general discounting of positive feedback and biased reasoning when evaluating one’s own abilities ^[2–5]^. Sufferers have been suggested to exhibit perfectionistic tendencies ^[6–8]^ and to over-prepare for tasks, believing they must always excel through sheer effort, since they doubt their natural abilities ^[9]^. This overwork can bring short-term success, but achievements are not internalised, and instead external factors (e.g. luck, timing) are credited ^[4,9]^.

Imposterism has been observed across various populations, educational settings and workplaces ^[10]^ and is associated with loss of motivation, reduced job satisfaction and overall deleterious consequences for an individual’s wellbeing ^[10–12]^. For instance, imposter feelings have been linked to symptoms of psychopathology, including anxiety and depression ^[13,14]^, low self-esteem ^[13,15]^ and social dysfunction ^[11]^. The apparent mis-calibration between self-confidence and objective achievements central to imposterism suggests that it may represent a deficit of metacognition, the process of being aware of and evaluating our own cognitive processes ^[16,17]^. Recent studies have shown that individuals high in imposterism report reduced confidence in their reasoning and problem-solving abilities relative to low-imposterism individuals, despite showing no objective decrement in performance ^[5,18]^. Previous studies of metacognition in imposterism have operationalised metacognition in terms of self-evaluation in tasks related to intelligence. However, it remains unknown whether metacognitive alterations extend beyond intellectual self-evaluation to other cognitive domains. It is possible that biased self-evaluation (low confidence) in imposterism may be restricted to specific sensitive domains (such as intelligence, education and/or work), or alternatively, low confidence might permeate metacognitive evaluation across most or all aspects of a person’s life (i.e., in a domain-general manner).

It remains unclear the degree to which confidence distortions in imposterism influence decision-making and behaviour. For instance, one crucial adaptive behaviour closely linked to confidence is information seeking. Confidence directly informs whether further information should be sought prior to decision commitment, with states of low confidence being associated with increased information-seeking behaviour regardless of initial accuracy ^[19–22]^. Persistent under-confidence in one’s abilities might lead to increased but inefficient and/or ineffective information-seeking behaviour. To our knowledge, this has not yet been investigated in imposterism.

Here, we investigated the relationships between imposterism, confidence and information seeking during performance of a perceptual decision-making task in a large general population sample (N = 740). This allowed us to assess whether confidence and information-seeking alterations associated with imposterism occur even during performance of a low-level perceptual task. We hypothesised that high imposterism would be associated with systematically low confidence (but not reduced accuracy) and increased (but ineffective/inefficient) information-seeking. Furthermore, by assessing perfectionism alongside imposterism, we sought to determine whether perfectionism may play a role in any observed relationships. Although the two constructs are strongly correlated ^[12,23,24]^, perfectionism reflects rigid self-standards and fear of mistakes ^[6]^, whereas imposterism involves discounting achievements and fears of fraudulence ^[1]^. Including both therefore allowed us to test whether effects on confidence and information-seeking are common to both traits or specific to imposterism.

## Materials and Methods

### Participants

740 participants were recruited overall (556 female/ 178 male/ 6 other, 17-80 years old (M = 40.16, SD = 13.87)). 43 participants were recruited using Sona Systems (https://www.sona-systems.com) and received undergraduate course credits for participating and 697 were recruited using Prolific (https://www.prolific.co) and were paid £6. Note that the participants in this study are a subset of those included in Mohr et al. (2024). The sample size was chosen to ensure adequate statistical power (80%) to detect effects at least as strong as those observed in a previous study from our lab ^[25]^. The power analysis indicated that 395 participants would be required to achieve 80% statistical power to detect such an effect. Hence, the large sample size allowed for high statistical power to be retained, even after data exclusion. All participants were offered a £20 reward if they achieved the top score on the information seeking task to motivate performance.

Pre-defined exclusion criteria (outlined below) led to the exclusion of 138 participants, leaving a final sample of 602 participants (458 female/ 144 male aged from 18 to 80 years (M = 41.20, SD = 13.89)). The overall exclusion rate (18.6%) is in line with previous similar online studies both from our own group ^[25]^ and others ^[22,26,27]^. The study received ethical approval from the University of Dundee Research Ethics Committee and University of Glasgow Ethics Committee (300210261). All participants provided informed consent.

### Perceptual Information-Seeking Task

Participants completed a two-alternative forced-choice (2-AFC) numerosity discrimination task. Figure 1 shows a schematic of the trial procedure. On each trial, a black cross appeared in the centre of a white screen for 1000 milliseconds (ms). This was followed by two black boxes, one on the left and one on the right side of the screen, which appeared simultaneously for 400 ms. Both boxes contained numerous white dots. The participant was asked to choose which box had contained a larger number of dots by pressing the ‘w’ key for left box or the ‘e’ key for right box. One box (the reference box) always contained 272 dots (out of 544 possible dot locations) while the other box contained an increased or reduced number of dots ranging from –64 to +64 dots (in increments of 8) in comparison to the reference. The location (left or right) of the reference box varied across trials and within each of the difficulty levels. The order of stimulus presentation was randomly generated for each participant within each block.

**Figure 1:**
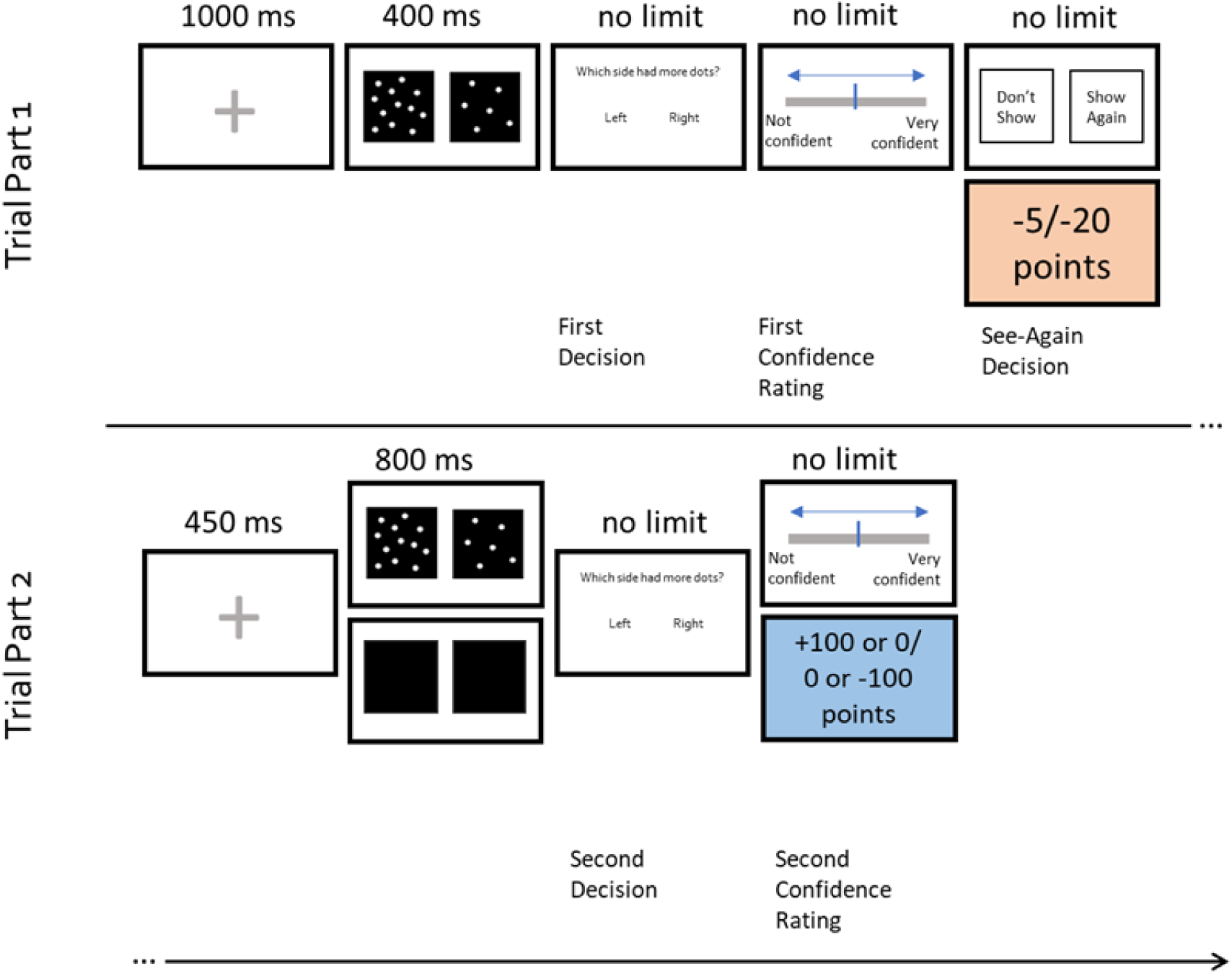
Information-seeking task. On each trial, participants judged which box (left or right) contained the higher number of dots and provided a confidence rating in each decision (scale of 1 to 6, where 1 represented “not confident (guessing)” and 6 represented “certain”). Participants were then given the chance to see the stimuli again for a certain cost and, depending on their choice, were subsequently shown either the stimuli or two blank black squares again. Participants were asked again to judge which box (left or right) contained the higher number of dots and provided a confidence rating. A correct final answer either provided a reward or the aversion of a loss. Participants were reminded of the cost of information seeking during the see-again decision on each trial but were not shown their score or points received/lost throughout the task.

Participants could respond after the stimulus screen changed to the response screen (i.e., not during the 400 ms of stimulus presentation). There was no time limit for the response and participants were not given feedback on whether their response was correct. After providing a response, participants were asked to rate ‘how confident are you that the decision was correct?’ on a scale of 1 (Not confident) to 6 (certain) using their mouse, equivalent to a half-confidence scale. Participants were instructed that if they were very unsure in their decision that they should choose the lowest (‘Not confident (guessing)’) option. There was no time limit for the confidence rating. To assess the tendency to seek further information in cases of uncertainty, after the first choice and confidence rating the participant was offered the chance to see the stimuli again (for a cost of either -5 or -20 points, depending on the block). If they chose to see the stimuli again, they were presented the same stimuli for a further 800 ms. Hence, seeing the stimuli again was always helpful. If they chose not to see again, they did not lose any points, but they instead saw two empty boxes for 800 ms. Regardless of whether they sought further information or not, the participant then made a final judgement on which box had contained more dots and provided a final confidence rating. To incentivise subjects to strive for the best possible overall accuracy, they either gained (appetitive blocks) or lost (aversive blocks) points based on their performance on this final decision. In appetitive blocks, if they were correct in this 2nd choice then they were awarded 100 points, whereas if they were wrong, they gained 0 points. In aversive blocks, if they were correct in the 2nd choice then they lost 0 points, whereas if they were wrong, they lost 100 points. Participants were reminded of the current rules (i.e., appetitive v aversive, low versus high information seeking cost) on each trial. Each participant completed a total of 128 trials (16 trials at each difficulty level, split evenly into 4 blocks consisting of 32 trials each). To counterbalance the order of information cost conditions, participants were randomly assigned to complete the experiment in either order A (appetitive/low-cost block; aversive/low-cost block; appetitive/high-cost block; aversive/high-cost block) or order B (appetitive/high-cost block; aversive/high-cost block; appetitive/low-cost block; aversive/low-cost block). Reward conditions always began with the appetitive block to enhance the sense of loss when previously earned points were later deducted for incorrect responses ^[28,29]^.

Participants could take self-paced breaks between blocks. Before starting the task, participants completed 4 practice trials in which they only had to make the initial decision (stimuli presentation, response, confidence rating) with feedback (a green tick for correct or a red cross for incorrect). Two further practice trials were used to familiarise participants with the confidence rating scale in which they were instructed how to respond if they were confident or not confident. Participants then completed 6 practice trials which had the same structure as the main task trials (stimuli presentation, response, confidence rating, see-again decision, stimuli presentation, response, confidence rating) but participants could see their cumulative score on the screen throughout these trials. Participants were not informed of their score at any point throughout the main task.

### Self-report questionnaires

Each participant completed the Clance Impostor Phenomenon Scale ^[3]^, and the Short Almost Perfect Scale ^[30]^. The inclusion of a questionnaire measuring perfectionism was motivated by its previously established link with imposterism ^[7,8,31]^. Perfectionism is thought to contribute to self-doubt and rigid self-evaluative tendencies in imposterism ^[8,12,24,32,33]^. Hence, we included a measure of perfectionism to establish whether it may play a role in any relationships between imposterism and the task measures.

Each participant also completed selected items (total = 63) from a battery of eight mental health questionnaires ^[34–42]^ which assessed symptomatology across a range of disorders. The items were identical to those used by ^[43]^ and allowed us to derive scores on an ‘Anxious-Depression’ (AD) symptom dimension ^[25–27,44]^. To calculate dimension scores for each participant, the raw responses for each item were first z-scored across participants, then the individual item z-scores within each participant were multiplied by their corresponding factor weights^[43]^ and the resulting products were summed across all items. Finally, the factor sums were z-scored across participants in preparation for statistical analyses. This allowed us to assess whether symptoms of anxiety and depression drive, or at least contribute to, any observed imposterism-behaviour relationships.

### Procedure

The experiment was conducted online via the Gorilla experiment platform ^[45]^ and could only be completed on either a laptop or computer (but not on a mobile phone or tablet). Participants were directed to an online consent form and information sheet. If informed consent was given, participants were asked to provide demographic information of age and gender. The participants then completed the questionnaires and task in a randomised order. The entire experimental session took approximately 1 hour.

### Exclusion Criteria

Several predefined exclusion criteria were applied to the data. Approximately 18.6% of participants were excluded, leaving 602 participants. Participants who met any one or more of the following criteria were excluded from all analyses:

- Did not provide gender information (n = 6, ∼1%).
- Below- or near-chance task performance (overall accuracy on first decision of the information-seeking task <55%) (n = 124, 16.8%).
- Used the same single confidence rating across all trials on the first decision of the task (n = 9, 1.2%).
- Incorrect response to a ‘catch’ item employed as an attention check (n = 8, 1.1%).

### Statistical analyses

#### Regression Models

We built a linear regression for each task measure (dependent variable) to examine the relationships with the survey measures, while controlling for age and gender:

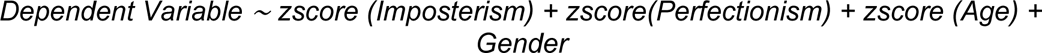

Continuous valued independent variables were z-scored prior to model fitting to obtain standardised regression coefficients. Gender was coded as female: -1, male: 1.

Additionally, to investigate the degree to which symptoms of anxious-depression (AD) contributed to observed relationships between imposterism and task measures, we performed additional hierarchical regression analyses. In each regression model, age, gender and AD scores were entered in the first step and imposterism scores were added in a second step to test whether adding imposterism as a predictor significantly increased the variance explained in the behaviour. The final models were as follows:

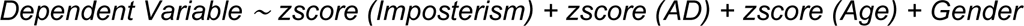

#### Multiple comparison correction

To address the problem of multiple comparisons, we controlled the false discovery rate (FDR) over the coefficients considered in each analysis using the Benjamini & Yekutieli (2001) procedure ^[47,48]^.

#### Mediation analysis

We conducted a single-level mediation analysis using the Mediation Toolbox (https://github.com/canlab/MediationToolbox) ^[49–51]^. Imposterism was entered as the predictor (X), confidence as the proposed mediator (M), and information-seeking as the outcome (Y). We first linearly regressed out age and gender from each variable and the mediation analysis was conducted on the residual values. All variables were subsequently z-scored prior to model estimation.

### Regression Analyses on Decision Change Metrics

To assess the relationships between imposterism and changes of mind (between the 1^st^ and final decision within-trials) as a function of whether further information was sought or not, we conducted a series of linear regression analyses using the following dependent variables:

- Number of changes in 1^st^-order response and confidence on information seeking trials.
- Number of changes in 1^st^-order response and confidence on non-seeking trials.
- Differences in accuracy and confidence between first and final decisions for seeking vs no-seeking trials
- Number of changes in 1^st^-order response and confidence following an initial correct response for seeking and non-seeking trials.
- Final accuracy and mean confidence following an initial correct response for seeking and non-seeking trials.

For this analysis, participants with extreme *information seeking* values were excluded, leaving 477 participants. Specifically, we removed any participants choosing to see the stimulus again on fewer than 5% or more than 95% of trials, thereby restricting the analysis to participants with sufficient trials sampled in both seeking and non-seeking conditions. Each model included imposterism, age, and gender as predictors. All variables were z-scored prior to analysis.

## Results

In an online study, participants from the general population (N = 602 after exclusion) performed an information-seeking task with confidence ratings (Figure 1) and completed the Clance Impostor Phenomenon Scale (CIPS), the Short Almost Perfect Scale (SAPS) and a battery of self-report psychiatric symptom questionnaires. The key task performance measures for each participant were first-order performance (indexed by Type-1 sensitivity (d’)) and confidence (indexed by average confidence ratings across trials) for both the first and final within-trial decisions separately. Additionally, we measured the percentage of trials in which participants sought additional information before making their second (final) decision. Sample distributions of CIPS scores, SAPS scores, tasks measures and demographics are shown in Supplementary Fig. S1.

### Adaptive use of information seeking in overall sample

As expected, participants were more likely to seek information on trials where they initially gave an incorrect response (Fig. 2A: t(601) = -27.6, p < .001), and final decision accuracy was higher for trials in which participants chose to seek information than for trials in which they chose not to (Fig. 2B: t(545) = 8.12, p < .001).

**Figure 2.**
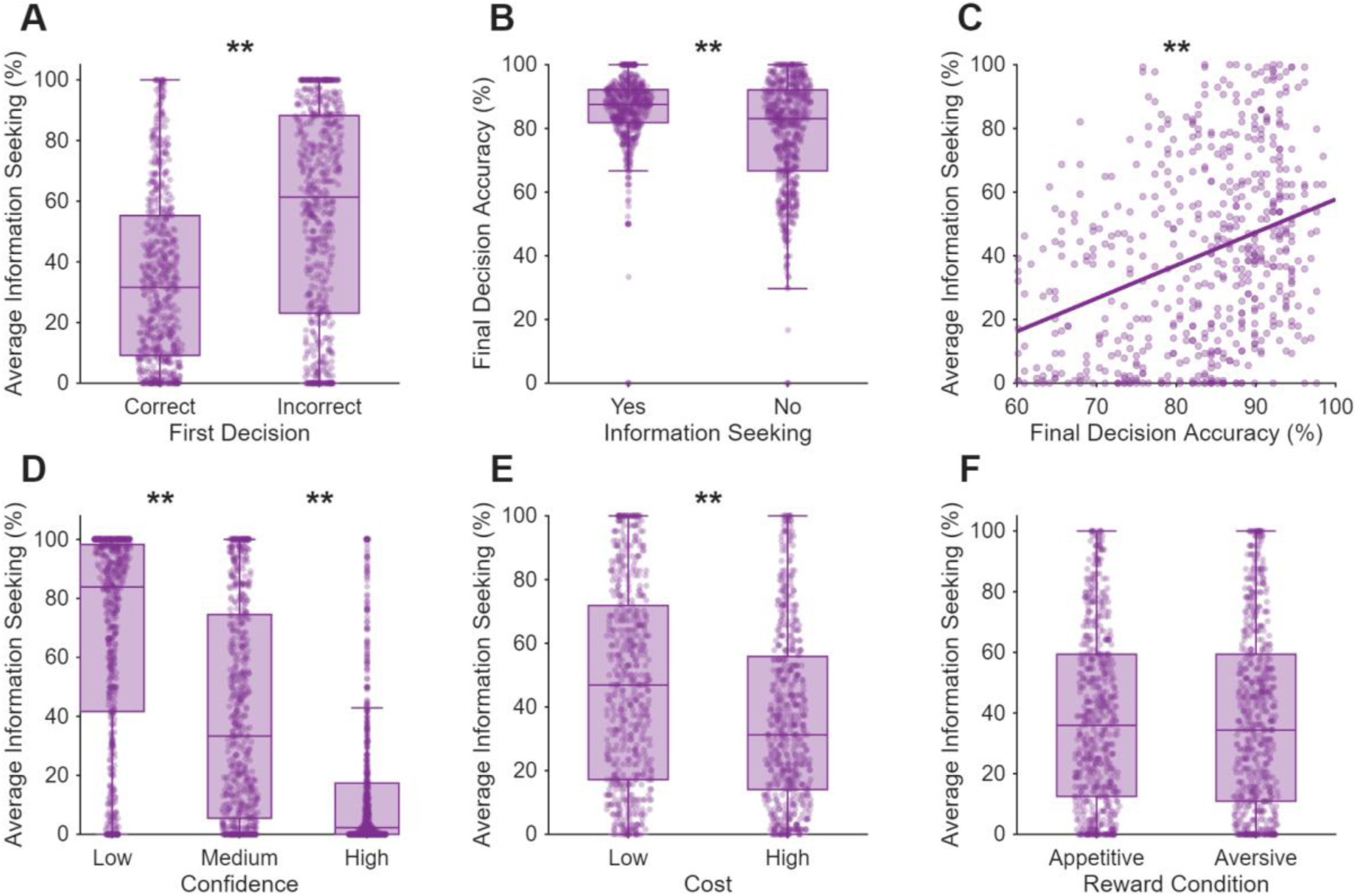
Overall use of information seeking. **A** Participants were more likely to seek information after an incorrect initial decision. **B** Participants were more likely to be correct in the final decision after choosing to see the additional information. **C** Participants who sought information more often had higher overall accuracy on the final decisions. **D** Participants used the information seeking option more when their initial decision confidence was lower. **E** Participants were more likely to seek information when the cost was low. **F** There was no difference in overall information seeking between appetitive and aversive trials. ** p <.001.

Between-participants, those who used the opportunity to seek information more often tended to be more accurate in their final decisions (Fig. 2C: r(600) = 0.38, p < .001). Additionally, participants were more likely to seek information when they reported low confidence in the initial decision (Fig. 2D: low versus medium confidence (t(589) = 23.7, p < .001), medium versus high confidence (t(595) = 23.4, p < .001)) and when the cost of seeking information was low (Fig. 2E: t(550) = 13.3, p < .001). Finally, participants were equally likely to seek information on appetitive and aversive trials (Fig. 2F: t(601) = 0.44, p = .66).

### Imposterism, but not perfectionism, is related to confidence and information seeking behaviour

To test our main hypotheses, we examined whether imposterism (CIPS scores) and/or perfectionism (SAPS scores) predicted task measures of accuracy, confidence and information-seeking. Figure 3 plots standardised regression coefficients indexing the strength and direction of the relationships between the survey scores and each task measure for both the 1st and final decisions, whilst accounting for age and gender (see Methods). We controlled the False Discovery Rate (FDR) at q < .05 over the 10 coefficients considered. As expected, imposterism and perfectionism were positively correlated (r(600) = .58, p <.001).

**Figure 3.**
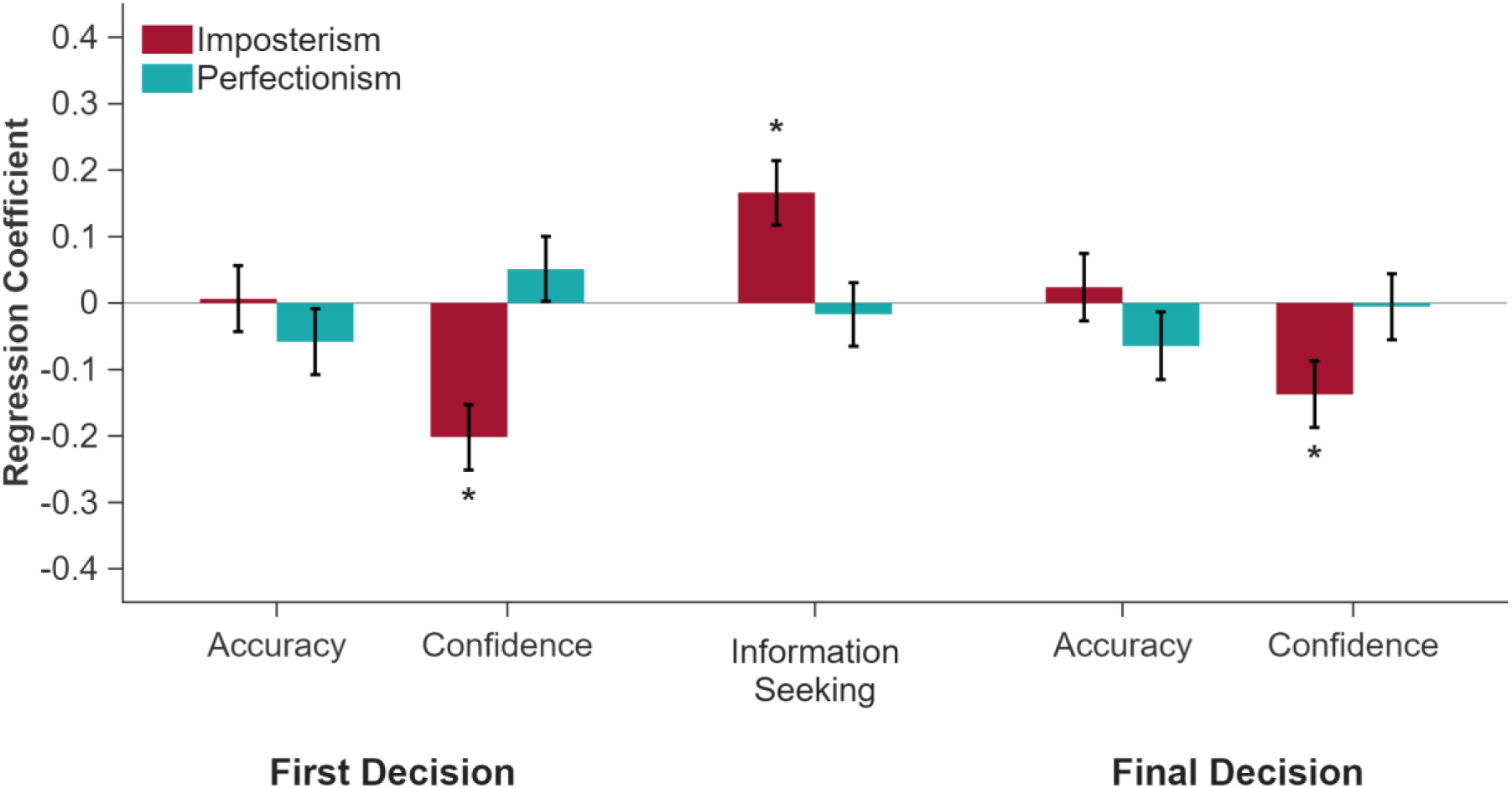
Relationships between behavioural measures and both imposterism and perfectionism, controlling for age and gender. Standardised regression coefficients (error bars show +/- standard error) from a linear model for each task measure (dependent variable) with age, gender (male: 1, female: -1) and questionnaire scores as independent variables. *q < 0.05 corrected for false discovery rate (FDR) over all coefficients.

In line with our hypotheses, higher imposterism was associated with lower average confidence in first decisions (β = –0.20, p < .001) but no reduction in objective accuracy (β = 0.006, p = .90). Consistent with low confidence in their initial choices, individuals high in imposterism were more likely to seek additional information (β = 0.16, p < .001) before committing to the final decisions. Intriguingly, despite being associated with increased information-seeking behaviour, high imposterism remained associated with low confidence in the final decisions (β = –0.14, p = .006) and still showed no significant relationship with objective accuracy (β = 0.02, p = .64). Hence, individuals reporting high levels of imposterism seemed to doubt their initial decisions and sought further information more often than individuals reporting low levels of imposterism, but this did not ameliorate their lower confidence or lead to increased objective performance. In contrast, perfectionism did not predict any of the task measures. It was not significantly related to first-decision confidence (β = 0.05, p = .30), first-decision accuracy (β = –0.06, p = .24), final-decision confidence (β = –0.01, p = .92), final-decision accuracy (β = – 0.06, p = .20), or information seeking (β = –0.02, p = .72).

Note that the imposterism-behaviour relationships were present independently of information cost and reward valence (see Supplementary Fig. S2). Additionally, the same pattern of results was observed when imposterism and perfectionism were entered separately (along with age and gender) into models predicting task measures (see Supplementary Fig. S3). We also conducted a series of control analyses to test whether the relationship between imposterism and information-seeking may be partially explained by inattentive responding^[52]^. The results did not support such an account (see Supplementary Results and Supplementary Fig. S4). No relationships were found between metacognitive sensitivity (meta-d’) or metacognitive efficiency (meta-d’/d’) and either imposterism or perfectionism (see Supplementary Fig. S5).

### Information-seeking behaviour dissociates imposterism from anxious-depression

To gain further insight into the nature of the relationships between imposterism and both confidence and information-seeking, we next investigated the role which symptoms of anxiety and depression may play. Imposterism is often associated with increased symptoms of anxiety and depression ^[13,14,53,54]^ and, indeed, in the current study imposterism scores were positively correlated with scores on an ‘anxious-depression’ (AD) transdiagnostic symptom dimension (r = 0.51, p < .001). Given that anxiety and depression have themselves been associated with low confidence in perceptual decisions ^[21,25,27,55]^, it is possible that AD symptoms may drive, or at least contribute to, the imposterism-behaviour relationships reported in Figure 3.

To test this, we carried out separate sequential hierarchical regression analyses for the three task outcome measures which showed significant relationships with imposterism (confidence in first decisions, information seeking and confidence in final decisions). In each regression model, age, gender and AD scores were entered in the first step and imposterism scores were added in a second step to test whether adding imposterism as a predictor significantly increased the variance explained in the behaviour. The full results of the hierarchical regression analyses are reported in Supplementary Table 1 and the final model regression coefficients for both AD and imposterism are shown in Figure 4A.

**Figure 4:**
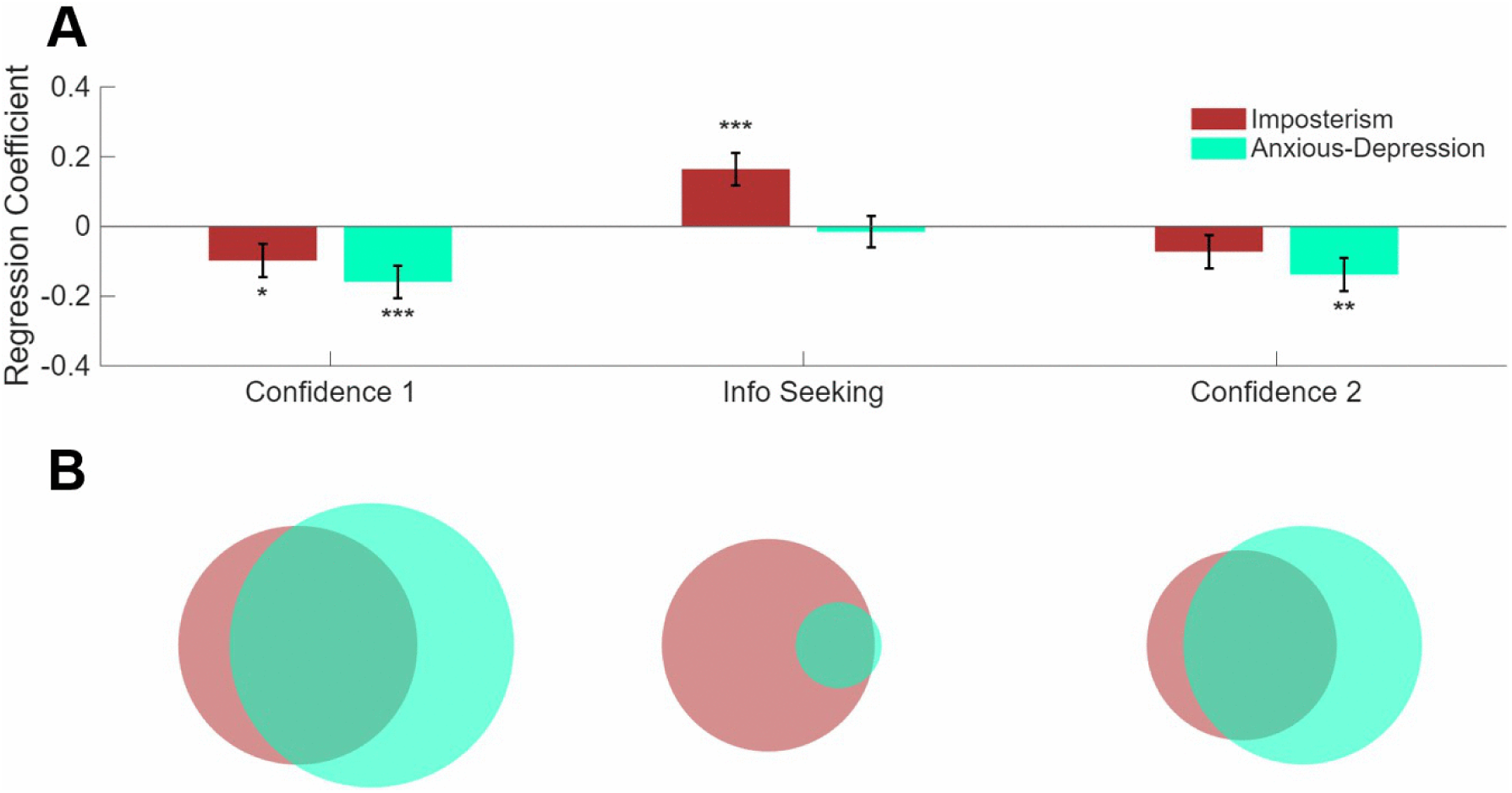
Shared and unique relationships between behavioural measures and both imposterism and anxious-depression. (A) Standardised regression coefficients (error bars show +/- standard error) from a linear model for each task measure (dependent variable) with age, gender (male: 1, female: -1), imposterism scores and Anxious-Depression scores as independent variables. *p < .05, **p < .01, ***p < .001. (B) Venn diagrams illustrating the total, shared and unique variance explained by imposterism and AD for each behavioural measure. Circle size is proportional to total R² explained by each predictor (over and above a baseline Age + Gender only model). The overlap represents the variance shared between the two predictors.

After controlling age and gender, AD symptoms were significantly negatively associated with confidence in first decisions (ΔR² = .04, ΔF(1,598) = 23.94, p < .001). However, the addition of imposterism to the model led to a significant increase in variance explained (ΔR² = .007, ΔF(1,597) = 4.372, p = .037). In the final model, both AD and imposterism were significant predictors. In contrast, AD symptoms showed no significant relationship with information seeking (ΔR² = .004, ΔF(1,598) = 2.377, p = .124) but again adding imposterism to the model led to a significant increase in variance explained (ΔR² = .019, ΔF(1,597) = 12.52, p < .001). In the final model, only imposterism significantly predicted information seeking. Finally, for confidence in final decisions, AD symptoms were significantly negatively associated (ΔR² = .028, ΔF(1,598) = 17.66, p < .001) and adding imposterism to the model did not significantly increase the variance explained (ΔR² = .004, ΔF(1,597) = 2.292, p = .131). In the final model, only AD significantly predicted confidence.

Figure 4B illustrates the total and shared variance explained (R²) by AD and imposterism on each outcome measure in the form of Venn diagrams. Overall, the results show that symptoms of anxiety and depression partially accounted for the relationship between imposterism and confidence, but imposterism uniquely predicted increased information seeking. Hence, information-seeking behaviour appears to distinguish between the overlapping yet partially distinct profiles of AD and imposterism.

### Local confidence partially mediates the relationship between imposterism and information-seeking

To formally test whether low first decision confidence explained the link between imposterism and increased information-seeking, we conducted a mediation analysis (Figure 5). We regressed out the effects of age and gender on all variables prior to the analysis. Higher imposterism scores predicted increased information-seeking (total effect: β = 0.16, p < .001) and this effect was partially mediated by low confidence in first decisions (mediation effect: β = 0.07, p < .001). However, the corrected direct relationship remained significant (β = 0.08, p = .025), indicating that a general tendency for increased information-seeking, which was independent of local confidence on the task at hand, also characterised imposterism.

**Figure 5.**
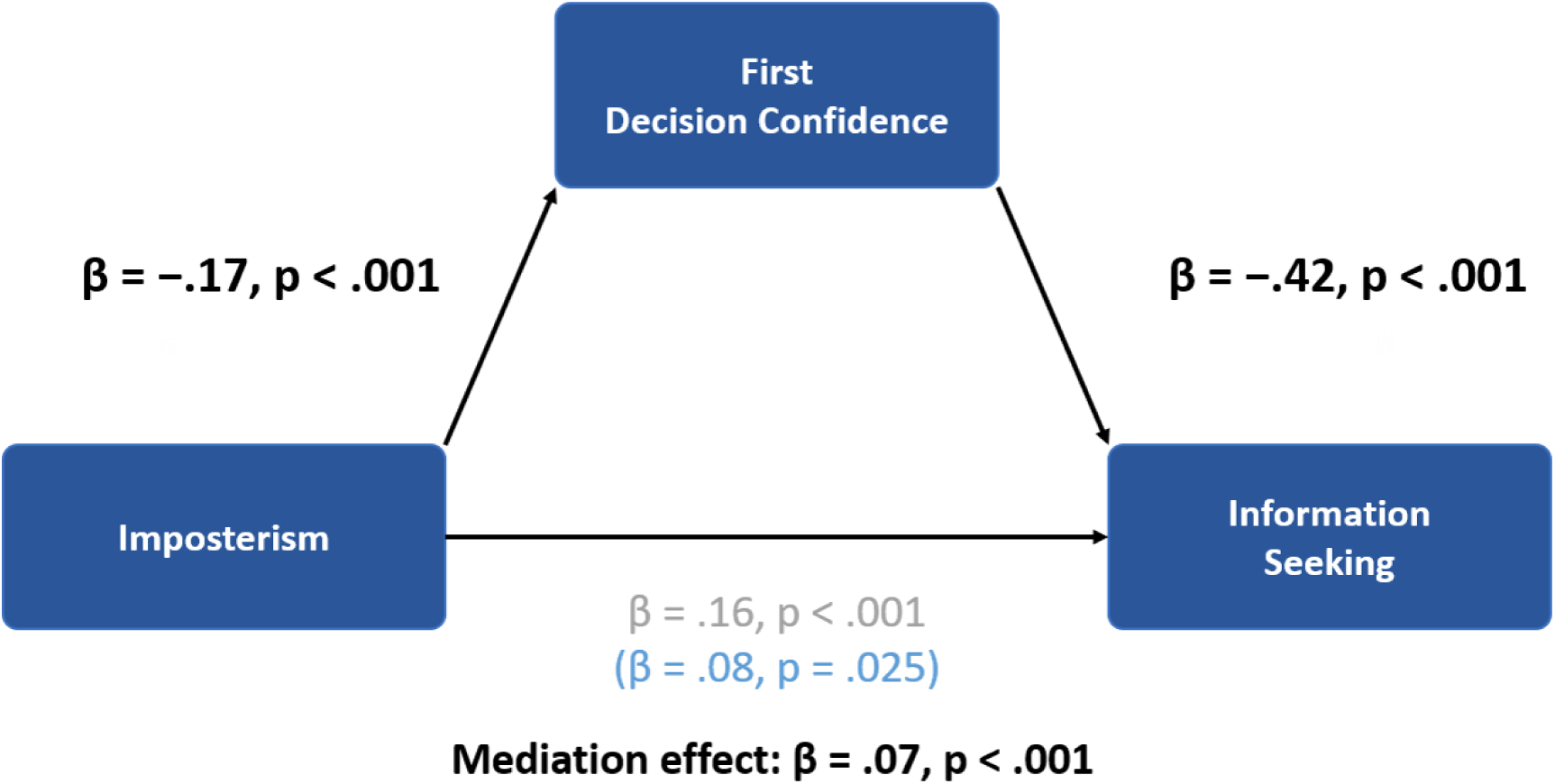
Mediation model (controlling for age and gender) showing that imposterism predicted greater information-seeking both directly and indirectly via reduced first decision confidence. The mediated relationship is shown in bold; the unmediated relationship is shown in blue, and the total relationship is shown in light grey.

### Increased information-seeking in imposterism compensates for maladaptive second-guessing

We were intrigued by the counterintuitive observation that despite seeking more information, high imposterism individuals remained low in confidence in their final decisions. Additionally, since in general information seeking was positively correlated with final decision accuracy in the full sample (r(600) = 0.38, p < .001) (see Figure S6), it is noteworthy that the imposterism-related increased information seeking did not lead to an imposterism-related increase in final decision accuracy. We posited two potential hypotheses to explain these observations.

First, seeing the stimuli again may have proven less helpful for high-imposterism participants and so, despite seeking more, their accuracy and confidence in the final decision did not change (**H1**: *ineffective seeking hypothesis*). For example, if high imposterism individuals did not incorporate the additional information appropriately into their final decisions. Under this hypothesis, imposterism should negatively correlate with changes in accuracy and/or confidence between the 1^st^ and final decisions on seeking trials only.

Alternatively, information seeking may have been equally helpful for both high and low imposterism individuals, but there may rather have been a difference in behaviour on trials in which they did not choose to seek information. Specifically, high-imposterism individuals may have differed in terms of changes-of-mind in non-seeking trials. For example, in the absence of additional information, the lower confidence in their 1^st^ decisions might have led to an increased tendency to change their response before the final decision, even if they were correct the first time. We labelled this hypothesis (**H2**) the *second-guessing hypothesis* because the increased information-seeking may have served to prevent maladaptive second-guessing which would otherwise result in lower performance in high-imposterism individuals.

To test these hypotheses, we regressed imposterism scores against changes in 1^st^-order performance and confidence between the 1^st^ and final decisions (controlling for age and gender), separately for trials in which participants chose to seek additional information and those in which they did not. Note that for these analyses, participants with extreme *information seeking* values (<5% or ≥95% of trials) were excluded, leaving 477 participants. The results are shown in Figure 6.

**Figure 6.**
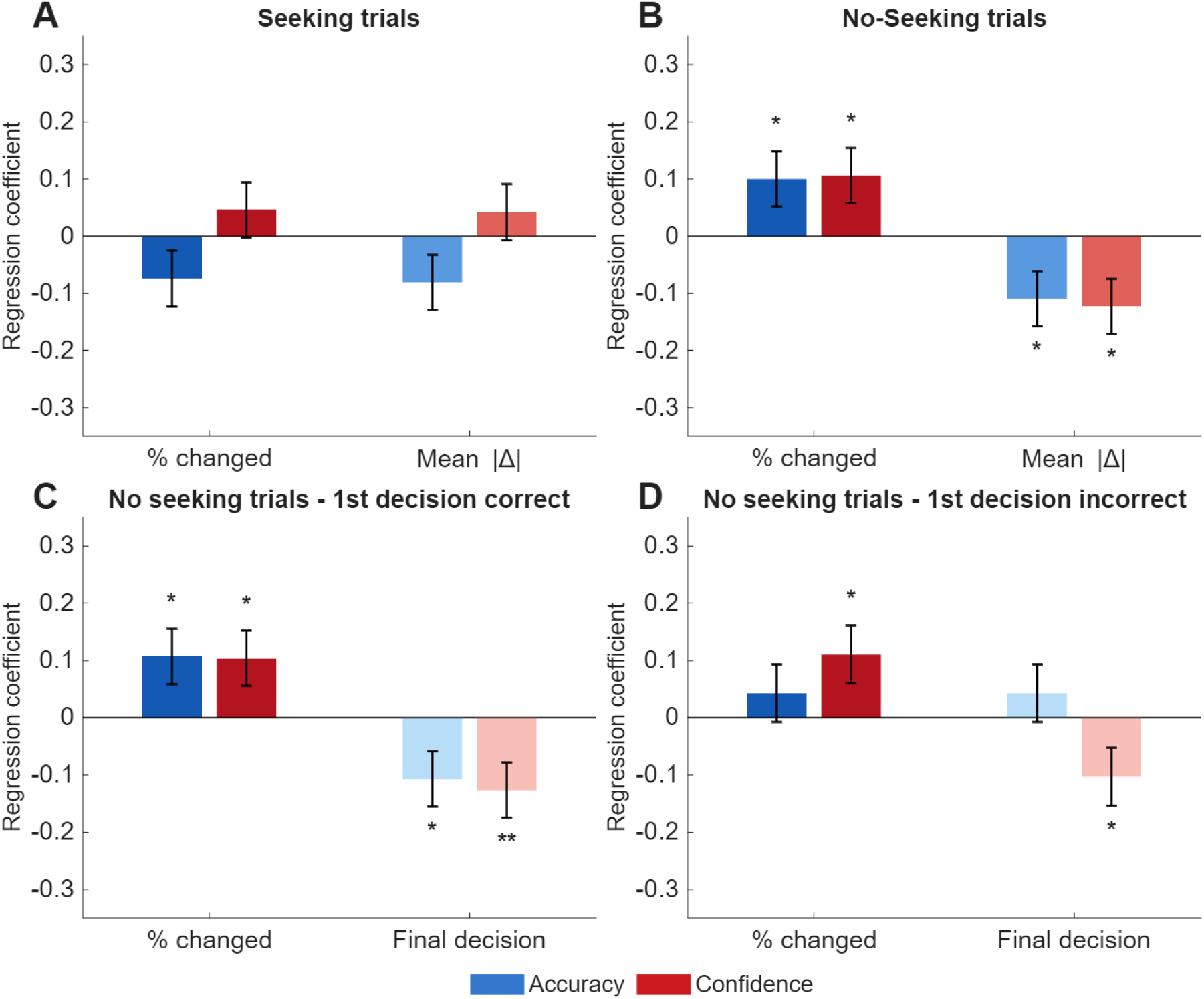
Relationships between imposterism and within-trial changes of mind for information seeking and non-seeking trials separately. Regression coefficients from linear models performed for each change-of-mind measure predicted by age, gender and imposterism, for trials in which participants did (A) and did not (B) seek information, and for non-seeking trials further separated by whether the first decision was correct (C) or incorrect (D). Error bars indicate standard errors of the coefficients and asterisks denote statistical significance (*p < .05, **p < .01).

For trials in which participants sought further information, imposterism was not significantly associated with the percentage of trials in which participants changed their initial response (β = –0.07, p = .13) or the percentage of trials in which they changed their initial confidence rating (β = 0.05, p = .34) (Fig. 6A). Likewise, imposterism did not significantly predict changes in decision accuracy (β = –0.08, p = .10) or confidence (β = 0.04, p = .38) between the first and final responses on seeking trials. These findings suggest that information seeking was effective for individuals across the imposterism spectrum, and do not support the notion that high-imposterism individuals derived less benefit from the additional information (**H1**: *ineffective seeking hypothesis*).

By contrast, the data provided support for the *second-guessing hypothesis (***H2**). On trials where participants *did not* seek additional information, higher imposterism was associated with significantly more changes in both 1^st^-order responses (β = 0.10, p = .04) and confidence ratings (β = 0.11, p = .03) between the first and final decisions. Crucially, in non-seeking trials (Fig. 6B), imposterism was associated with a reduction in accuracy (β = –0.11, p = .02) between first and final decisions, indicating a tendency to switch more often from correct to incorrect responses. Imposterism also predicted larger reductions in confidence (β = –0.12, p = .01), suggesting increased uncertainty in the absence of external reassurance. To further unpack these effects, we separated the non-seeking trials into those in which participants were correct in their first decision (Fig. 6C) and those in which they were incorrect (Fig. 6D). When first decisions were correct, higher imposterism was associated with significantly more response (β = 0.11, p = .03) and confidence (β = 0.10, p = .03) changes from the first to the final decisions, which resulted in reductions in both final accuracy (β = –0.11, p = .03) and confidence (β = – 0.13, p = .008). Hence, when they did not seek information, individuals high in imposterism were more likely to abandon a correct initial answer — switching away from it and becoming less certain in the process. By contrast, when the first decision was incorrect, imposterism was not significantly associated with response (β = 0.04, p = .40) or accuracy (β = 0.04, p = .40) changes, though it did still predict increased confidence changes (β = 0.11, p = .03) which resulted in lower final decision confidence (β = -0.11, p = .03). This confirms that the increased changes of mind observed in non-seeking trials reflect maladaptive second-guessing rather than adaptive error correction, providing support for H2. Together, our results indicate that the cognitive profile of high-imposterism individuals is characterised not by ineffective use of information, but by a heightened vulnerability to second-guessing in its absence.

## Discussion

The current study provides robust evidence that imposterism is associated with alterations in confidence and information-seeking behaviour during perceptual decision-making. In a large general population sample, we found that individuals with higher imposter traits exhibited consistently low confidence in their perceptual decisions, despite exhibiting no deficits in objective performance. High-imposter individuals also demonstrated an increased tendency to seek additional information before committing to final decisions. This increased information-seeking was partly mediated by low initial decision confidence and served to buffer against a maladaptive tendency for high-imposterism individuals to second-guess themselves in the absence of new evidence, by switching from a correct to an incorrect answer. Notably, these behavioural patterns were not associated with perfectionism, despite the two traits being moderately to strongly correlated. Together, these findings delineate specific metacognitive signatures of imposterism, highlighting its influence on self-monitoring and decision regulation.

### Imposterism and decision confidence

The negative correlation between imposterism and decision confidence converges with prior research linking imposter feelings to low self-confidence in academic and professional domains ^[1,4,10–13,15,33]^. Here, we demonstrate that this under-confidence generalises to perceptual decision-making contexts devoid of social or intellectual evaluation, and hence likely reflects a pervasive metacognitive bias not restricted to sensitive self-domains. Importantly, the under-confidence occurred in the absence of any reduction in decision accuracy, underscoring a dissociation between subjective confidence and objective ability in high-imposterism individuals. This is consistent with models positing that imposter feelings arise not from actual incompetence, but from maladaptive internal attribution styles and distorted self-evaluation ^[1,2,4,13]^. From a metacognitive perspective, it suggests a disrupted calibration between confidence and performance, potentially implicating impairments in error monitoring and/or self-referential evaluation mechanisms ^[55]^.

### Information-seeking compensates for maladaptive second-guessing

Individuals higher in imposter traits were more likely to opt to view the stimulus again before making a final decision. Mediation analysis revealed that this increased information-seeking behaviour was partially explained by low confidence in initial decisions, consistent with models in which information-seeking serves as a strategy to reduce uncertainty or bolster confidence ^[19–21]^. However, a significant direct effect of imposterism on information-seeking remained even after accounting for initial confidence, indicating that this behaviour also reflects a more generalised need for reassurance or verification among high-imposterism individuals which occurs regardless of their estimation of ‘local’ performance on the task-at-hand. One plausible hypothesis is that the direct effect of imposterism on information-seeking may arise from an imposterism-related reduction in overall ‘global’ self-beliefs ^[44,57]^. In other words, high-imposterism individuals may generally assume that they are likely to be bad at a given task, independently of any direct evaluation of their actual performance, and this assumption may also influence their information-seeking behaviour. Further studies incorporating measures of ‘global’ confidence can test this hypothesis.

Interestingly, increased information-seeking did not lead to corresponding gains in accuracy or final decision confidence in high-relative to low-imposter individuals. This might initially suggest that the additional information gained by high imposter individuals was not effectively integrated into the decision process. However, follow-up analyses did not find evidence to support this interpretation. On trials where participants chose to seek additional information, imposterism was not associated with a difference in how often they changed their minds, nor did it relate to changes in accuracy or confidence between first and final decisions. In other words, both high- and low-imposterism individuals seemed to derive similar objective benefit from information-seeking, indicating that the use of information was not impaired.

Instead, we observed clear evidence supporting a “second-guessing” hypothesis. On trials where participants did not seek additional information, individuals high in imposterism were more likely to change their answers, even when they were initially correct, and more likely to reduce their confidence between initial and final decisions. This tendency to second-guess oneself in the absence of new evidence was maladaptive, as it was associated with decreased accuracy. These findings align with characterisations of imposterism as a state marked by excessive doubt, rumination, and hypervigilance towards perceived failure^[7,8,31]^. In our data, the elevated information-seeking observed among high-imposterism individuals appeared to serve as a protective mechanism which offset, to some extent, a metacognitive instability that otherwise undermined performance.

### Dissociating imposterism from perfectionism

A notable feature of our results was the absence of any behavioural effects associated with perfectionism, despite its moderate correlation with imposterism at the trait level. It has recently been proposed that cognitive-behavioural differences can provide invaluable insights when attempting to dissociate over-lapping but distinct dimensions of personality/psychopathology ^[58]^ and our results support this. Perfectionism has long been considered a core component, or even a driver, of the imposter phenomenon ^[1,7,8,31]^, and both constructs are associated with maladaptive evaluative concerns and fear of failure ^[12,23,24]^. However, perfectionism in our study did not predict task confidence or information-seeking behaviour, whilst imposterism did. This dissociation is theoretically important. It suggests that while perfectionism and imposterism may co-occur, they impact cognition and decision-making in distinct ways.

One possible interpretation is that perfectionism primarily influences goal setting and standards, whereas imposterism exerts stronger effects on self-evaluation and confidence calibration. For example, a perfectionist might experience distress at the prospect of imperfection but still maintain a coherent and accurate appraisal of their abilities. In contrast, imposterism appears to manifest as a more pervasive distrust in one’s own competence, even in the face of objective success. This distinction may have critical implications for intervention design. Whereas perfectionism might respond to strategies targeting unhelpful standards (e.g., cognitive restructuring ^[59]^), imposterism may require interventions focused on metacognitive awareness ^[60,61]^, attributional style ^[62]^, and/or self-efficacy enhancement ^[63]^. Building on this, a related interpretation is that while the two constructs are closely intertwined, perfectionism may centre on anticipatory evaluation of future performance, whereas imposterism relates more to retrospective evaluation of past performance. This aligns with our data, but further work would be needed with new experimental designs to determine if perfectionism can be linked to anticipatory evaluations of upcoming behaviours.

### Overlap and divergence between imposterism and a transdiagnostic anxious-depression dimension

The results refine our understanding of the cognitive processes associated with imposterism by demonstrating both considerable overlap with, but also subtle distinctions from, symptoms of anxiety and depression. In line with previous studies, imposterism was positively associated with anxious-depression symptoms ^[53,54]^ and individuals reporting high levels of anxiety and depression also reported systematic low confidence in perceptual decisions, despite showing no detriment in objective accuracy ^[21,25,27,55]^. However, hierarchical regression analyses revealed that imposterism contributed additional explanatory power to initial decision confidence beyond that accounted for by anxiety and depression. More strikingly, information-seeking behaviour emerged as a distinctive feature of imposterism. Whereas anxious-depression showed no significant association with information seeking, imposterism remained a significant predictor even after controlling for age, gender, and anxious-depression symptoms. In contrast, confidence in final decisions was primarily associated with anxious-depression symptoms, implying that broader affective factors may exert a more pervasive influence on subjective confidence even after uncertainty has been reduced through additional information gathering. Importantly, these findings should not be interpreted as demonstrating that imposterism is fully distinct from anxious-depression. Rather, the present results indicate substantial overlap between the constructs, while suggesting that increased information-seeking may capture variance associated with imposterism that is not explained by anxious-depression symptoms. Future work including wider behavioural measures, potentially in longitudinal designs combined with latent variable approaches, would help clarify the extent to which imposterism reflects a construct that is separable from anxious-depression.

### Theoretical and practical implications

By highlighting reduced metacognitive confidence and increased reliance on reassurance-seeking strategies, this study supports models conceptualising imposterism as a metacognitive disorder: a failure not in competence per se, but in the regulation and appraisal of competence ^[16,64]^. This perspective aligns with broader theories of anxiety and self-doubt in which metacognitive dysregulation plays a central role ^[61,65]^.

Practically, the results offer insights for clinical, educational, and organisational interventions. Imposter feelings are common among high-achieving individuals and have been linked to burnout, reduced job satisfaction, and career disengagement ^[10–12]^. If, as our data suggest, these feelings stem from pervasive under-confidence and a tendency towards maladaptive second-guessing, then interventions aimed at improving metacognitive calibration and promoting adaptive self-evaluation may be particularly beneficial. Techniques such as confidence training ^[66]^ and/or metacognitive therapy ^[61]^ may hold promise in this regard.

Our findings also raise intriguing possibilities for the design of decision-support systems or workplace feedback mechanisms. For example, structured information-seeking opportunities or confidence tracking tools might help high-imposterism individuals better calibrate their decisions and reduce unnecessary cognitive load. However, such tools should be carefully designed to foster autonomy rather than dependency on external reassurance.

### Methodological strengths and limitations

The robustness of our findings is underscored by several methodological strengths. First, we employed a large, diverse sample (N = 602), providing high statistical power. Second, the core results were consistent across multiple task conditions (separate trial subsets), mitigating concerns about overfitting or false positives ^[67]^. Third, by controlling for age and gender, we were able to isolate specific associations between trait measures and task behaviour.

However, some limitations should also be acknowledged. The study relied on self-report measures of imposterism and perfectionism, which are susceptible to biases such as social desirability or limited introspective access ^[68]^. Future research could complement these with behavioural or clinician-rated assessments. Additionally, because the task was designed to capture confidence and information-seeking behaviours in an online setting, it necessarily abstracts away from the complex social and professional contexts in which imposter feelings have typically been reported to arise. It will be important for future studies to investigate whether similar (meta)cognitive patterns emerge in more naturalistic or high-stakes settings, such as academic evaluations or job performance reviews.

## Conclusion

In sum, we have shown that imposterism is associated with pervasive low confidence and increased information-seeking during perceptual decision-making. The increase in information-seeking compensated for an imposterism linked tendency to maladaptively second-guess oneself in the absence of additional information. Despite their objective competence, high-imposterism individuals display systematic disruptions in self-monitoring and confidence regulation, which may underlie the emotional and functional impairments associated with the imposter phenomenon. By identifying specific (meta)cognitive and behavioural signatures of imposterism, our findings open the door to targeted interventions and provide a foundation for future work on the mechanisms and consequences of this widespread psychological experience.

## Data Availability

All data are openly available on the Open Science Framework (OSF) under the URL: https://osf.io/s7kfx.

## Acknowledgments

This study was supported by the British Academy/Leverhulme Trust and the United Kingdom Department for Business, Energy and Industrial Strategy [SRG19/191169] and the United Kingdom Engineering and Physical Sciences Research Council (EPSRC) [EP/R513222/1].

