## Supplementary Material for "The metacognitive signatures of imposterism"

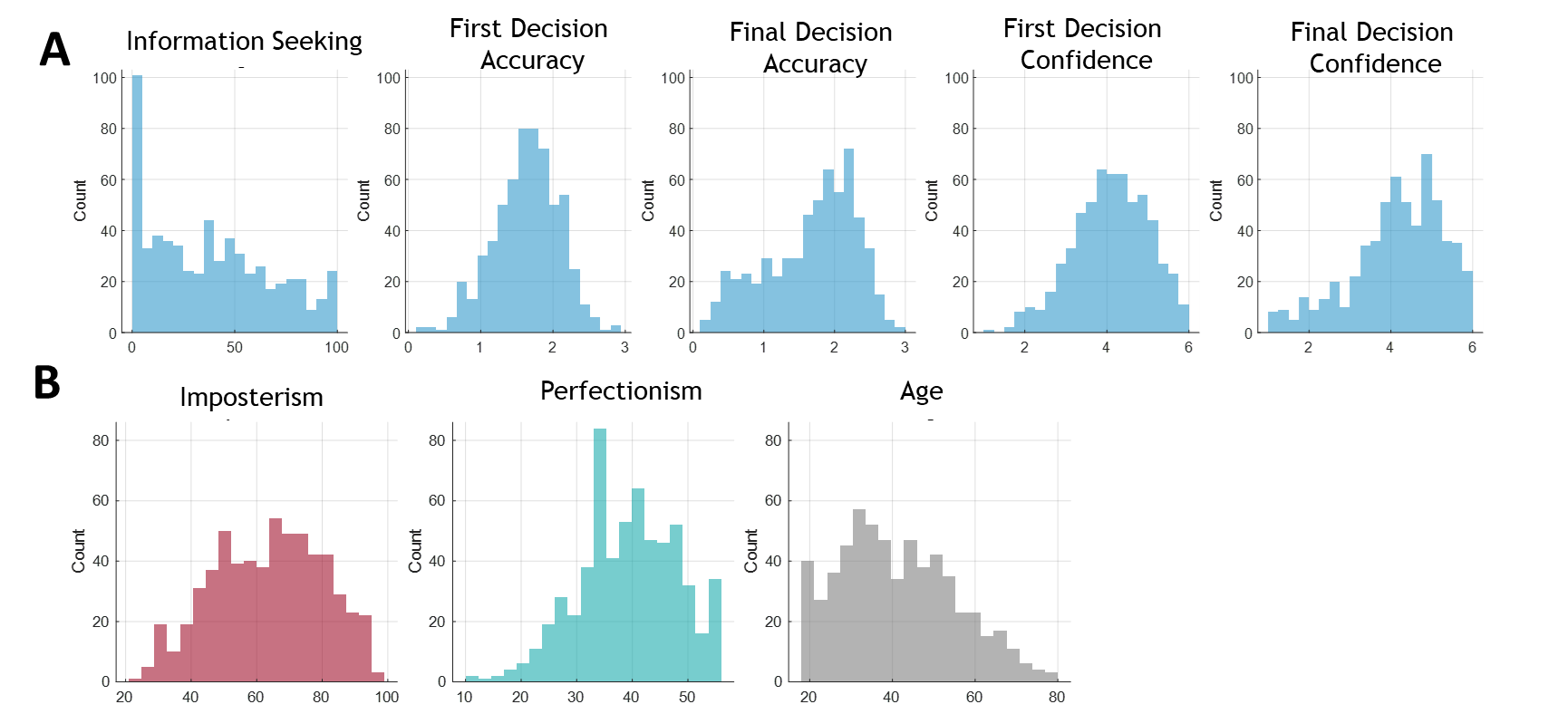

Figure S1. Distributions of key variables across the sample (N = 602). Top row: behavioural measures from the perceptual information-seeking task, including proportion of information-seeking trials. accuracy (first and final decisions), and confidence ratings (first and final decisions). Bottom row: self-report measures and demographics: Clance Impostor Phenomenon Scale (CIPS) scores (red), Short Almost Perfect Scale (SAPS) scores (blue), and participant age (grey).

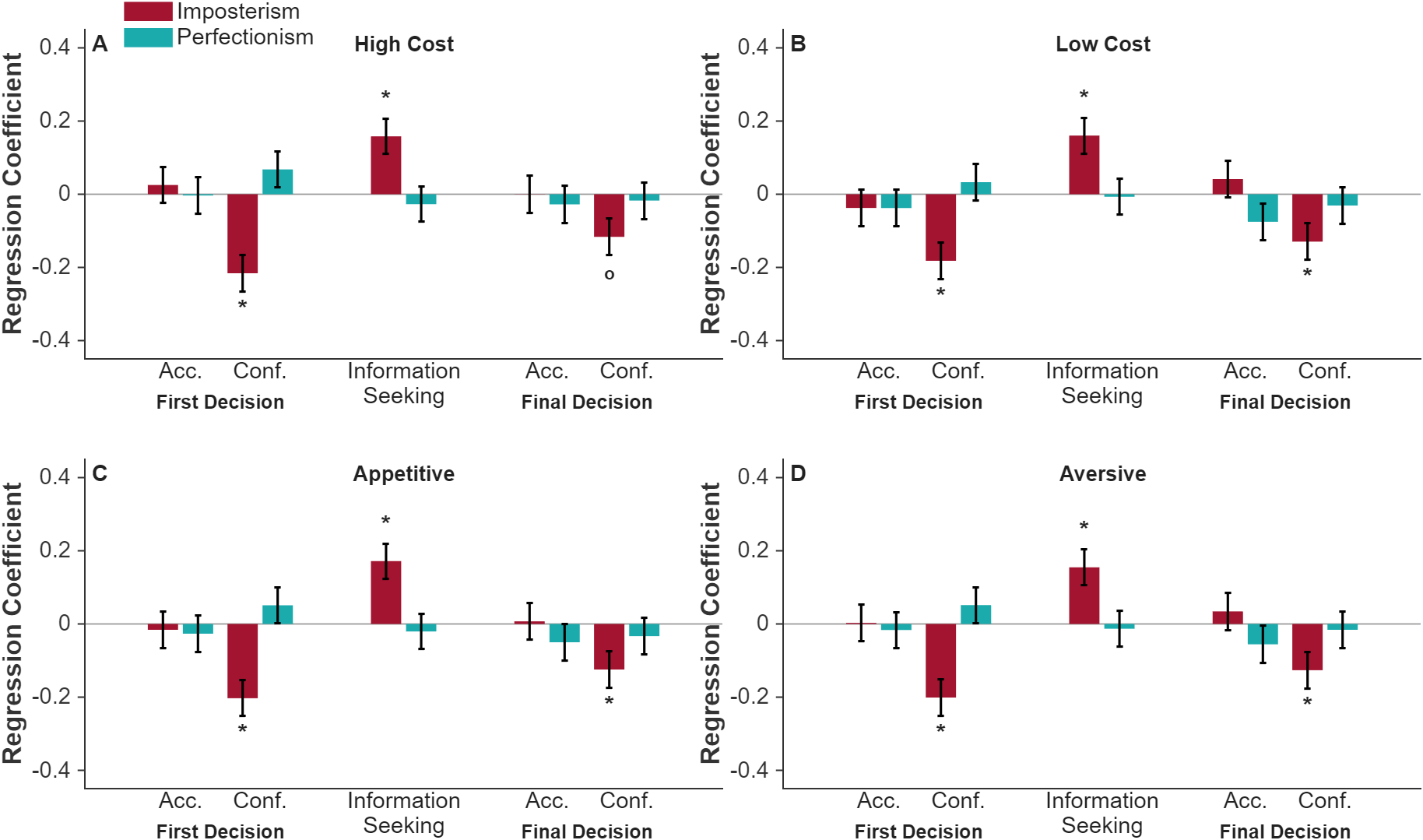

Figure S2. Standardised regression coefficients show the relationships between task measures and both imposterism and perfectionism, controlling age and gender, across reward and cost conditions. Error bars represent ±1 standard error. Asterisks indicate coefficients surviving false discovery rate (FDR) correction at q < .05.

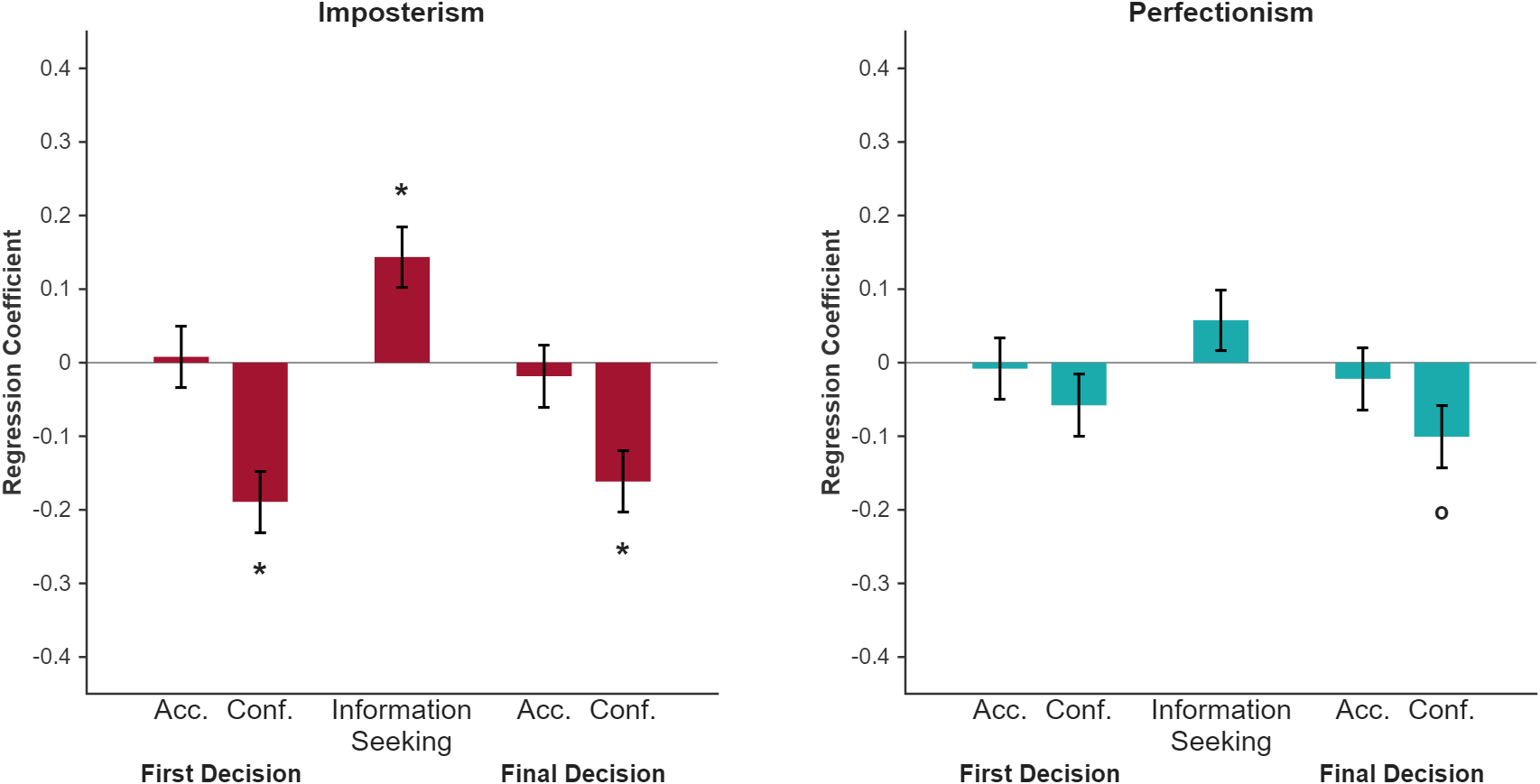

Figure S3. Standardised regression coefficients (± SE) showing the relationships between task measures and both imposterism (left) and perfectionism (right) separately. All models controlled for age and gender. * indicates FDR-corrected significance; o indicates p < .05 (uncorrected).

Supplementary Results

Robustness of results to inattentive responding

It has recently been shown have shown that inattentive responding can contribute to spurious relationships being detected between self-reported questionnaire scores and task measures (Zorowitz et al., 2023). Of relevance to the current study, if inattentive responders scored extreme values on the imposterism scale AND happened to never (or rarely) seek information, then this could lead to a relationship being observed between imposterism and information-seeking which was primarily driven by inattentive responding. Hence, we investigated the likelihood that inattentive responding may be contributing to the observed imposterism-information seeking relationship.

First, we plotted scores on both the imposterism and perfectionism questionnaires as a function of seeking behaviour (<5% seeking trials versus ≥5% seeking trials):

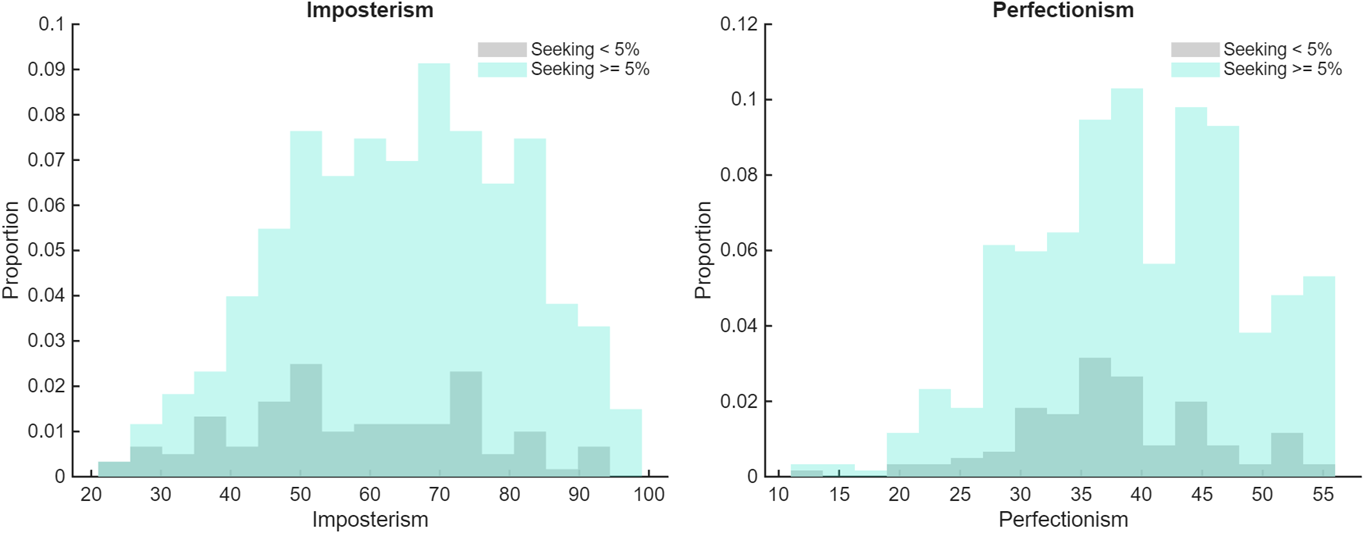

Figure S4. Distribution of Imposterism Scale scores and Perfectionism Scale scores stratified by information-seeking behaviour (seeking on <5% of trials vs seeking on ≥5% of trials). Histogram bars represent proportions of the full sample.

According to Zorowitz et al., (2023), positively skewed symptom scores are a particular problem because most people in the general population report few symptoms, so the “true” distribution is heavily weighted toward low scores. That means inattentive/careless responders are more likely to look artificially symptomatic (inflated scores), which can create spurious correlations between symptom severity and task performance rather than reflecting real effects. It is notable that neither the imposterism nor perfectionism questionnaires here showed right-skewed distributions and non-seekers did not show inflated scores compared to seekers. Indeed, independent samples t-tests revealed that non-seekers scored significantly lower for imposterism than seekers (t(600) = -3.6, p < .001) in line with the negative relationship between imposterism and information seeking observed in our main analyses. Non-seekers also scored lower than seekers for perfectionism, but the effect was not statistically significant (t(600) = -1.9, p = .052).

The information seeking measure itself is not an appropriate index of inattentive responding because not seeking information can be a valid strategy to maximise points on the task if one is confident in their 1st decisions. Importantly, choosing to see the stimulus again or not did not change the overall length of each trial (and therefore the overall experiment), a design choice we made to try to prevent the incentive for careless participants to rush through the experiment by not seeking information.

More appropriate indices of inattentive responding are provided by choice accuracy (operationalised as the primary task index of inattention by Zorowitz et al.) and overall points accumulated on the task. If inattentive responding characterised low imposterism participants, we would have expected imposterism to correlate positively with task accuracy (for the first and/or final decisions within trials) and/or total points accumulated on the task. However, imposterism was not significantly related to either task accuracy (first decisions: β = 0.006, *p* = .90; final decisions: β = 0.02, *p* = .64) or points accumulated on the task (β = −0.06, *p* = .16). However, to more directly test whether inattention might have played a role in the imposterism–information-seeking relationship, we carried out an additional multiple regression analysis with age, gender, imposterism scores and first-decision accuracy entered as predictors of information seeking. If inattentive responding was a confounder, we would expect that controlling for accuracy in the regression model would weaken or abolish the relationship between imposterism scores and information seeking. However, this was not the case. Although first-decision accuracy significantly predicted information-seeking behaviour (β = 0.163, *p* < .001), it did so independently of imposterism, which also significantly predicted information seeking (β = 0.160, *p* < .001). Indeed, including accuracy in the model slightly increased, rather than weakened, the association between imposterism and information seeking (β = 0.157 without accuracy; β = 0.160 with accuracy). Hence, we found no empirical evidence that inattentive responding was driving the imposterism–information-seeking relationship.

Finally, we note that Zorowitz et al., (2023) found that excluding participants who failed an attention check item embedded in the questionnaires almost entirely abolished the spurious relationships they observed. In the current study, we embedded a ‘catch’ item (“If you are paying attention, please select ‘Good part of the time’ for this answer”) within the questionnaire battery and excluded all participants who did not select the correct response (n = 8 excluded, 1.1% of full sample). Additionally, we excluded all participants who failed to achieve above-chance task performance (overall accuracy on first decision of the information-seeking task ≥55%) (n = 124 excluded, 16.8% of full sample). We are hopeful that these controls largely mitigated against the possibility of inattentive responding confounding the results.

Altogether, we are confident that inattentive/careless responding is unlikely to confound the relationship between imposterism and information seeking observed in the current study.

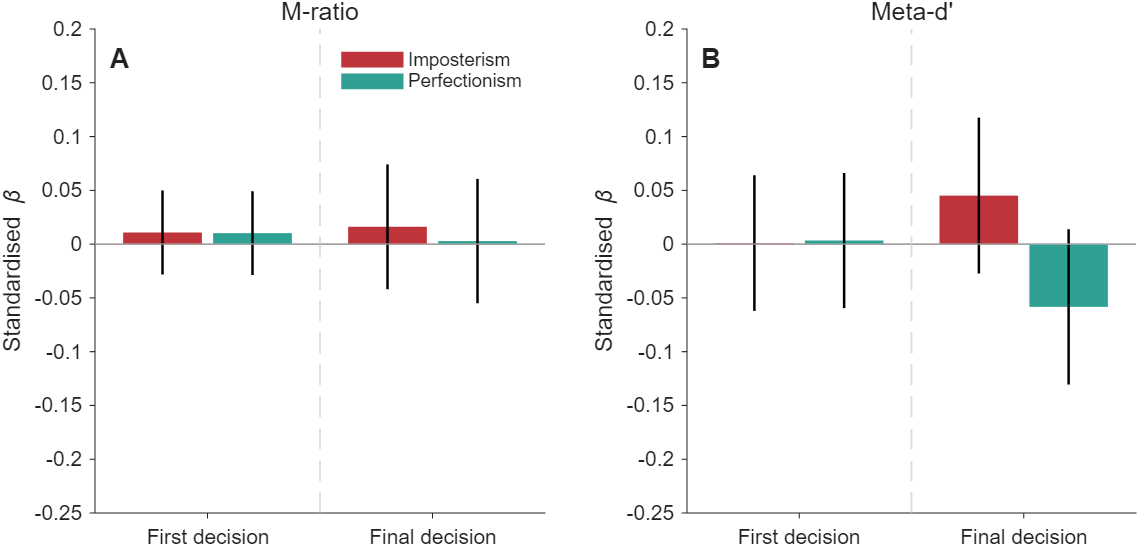

Figure S5. Relationships between metacognitive ability and both imposterism and perfectionism, controlling age and gender. Standardised regression coefficients (β) with standard error bars for **(A)** M-ratio and **(B)** meta-d' at first and final decisions.

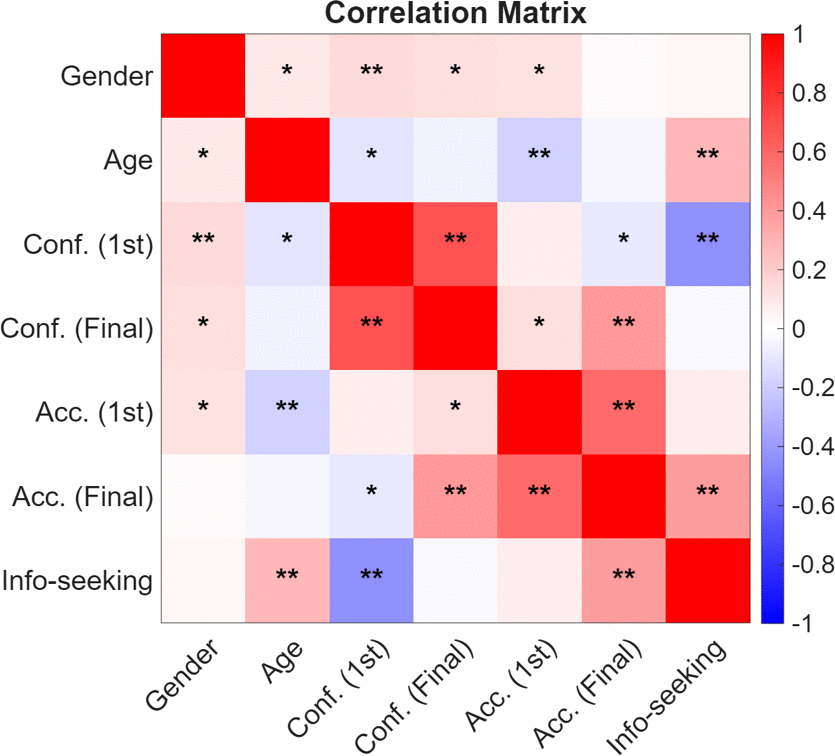

Figure S6. Correlation matrix of task and demographic variables. Pairwise Pearson correlations are shown between task measures (confidence, accuracy, information-seeking) and demographic variables (age, gender; gender coded 1 = male, -1 = female). Asterisks mark significant correlations (*p < .05, **p < .001).

**Table S1.** Hierarchical regression analyses predicting confidence and seeking behaviour.

|  | **Model** | **Predictor** | ***β*** | ***p*** | **Model *R*^2^** |
| --- | --- | --- | --- | --- | --- |
| ***Confidence 1*** | Model 1: Age + Gender | Age | −0.121 | .003 | .036 |
|  |  | Gender | 0.185 | < .001 |  |
|  | Model 2: + CIPS | Age | −0.170 | < .001 | .065 |
|  |  | Gender | 0.165 | < .001 |  |
|  |  | CIPS | −0.176 | < .001 |  |
|  | Model 3: + AD | Age | −0.165 | < .001 | .076 |
|  |  | Gender | 0.179 | < .001 |  |
|  |  | AD | −0.205 | < .001 |  |
|  | Model 4: + CIPS + AD | Age | −0.183 | < .001 | .083 |
|  |  | Gender | 0.169 | < .001 |  |
|  |  | CIPS | −0.098 | .037 |  |
|  |  | AD | −0.159 | < .001 |  |
| ***Information seeking*** | Model 1: Age + Gender | Age | 0.273 | < .001 | .075 |
|  |  | Gender | 0.010 | .821 |  |
|  | Model 2: + CIPS | Age | 0.317 | < .001 | .098 |
|  |  | Gender | 0.028 | .547 |  |
|  |  | CIPS | 0.157 | < .001 |  |
|  | Model 3: + AD | Age | 0.287 | < .001 | .079 |
|  |  | Gender | 0.012 | .792 |  |
|  |  | AD | 0.062 | .124 |  |
|  | Model 4: + CIPS + AD | Age | 0.316 | < .001 | .098 |
|  |  | Gender | 0.028 | .541 |  |
|  |  | CIPS | 0.165 | < .001 |  |
|  |  | AD | −0.015 | .735 |  |
| ***Confidence 2*** | Model 1: Age + Gender | Age | −0.054 | .188 | .017 |
|  |  | Gender | 0.145 | .002 |  |
|  | Model 2: + CIPS | Age | −0.093 | .028 | .035 |
|  |  | Gender | 0.129 | .007 |  |
|  |  | CIPS | −0.140 | .001 |  |
|  | Model 3: + AD | Age | −0.090 | .028 | .045 |
|  |  | Gender | 0.140 | .003 |  |
|  |  | AD | −0.172 | < .001 |  |
|  | Model 4: + CIPS + AD | Age | −0.103 | .014 | .049 |
|  |  | Gender | 0.133 | .005 |  |
|  |  | CIPS | −0.072 | .131 |  |
|  |  | AD | −0.138 | .003 |  |

*Note.* CIPS = Clance Impostor Phenomenon Scale; AD = Anxious-Depression. Standardised regression coefficients (*β*) are reported.
